# Spatially resolved analysis of growth dynamics in pome and drupe fruits of Rosaceae using 3D Gaussian Splatting

**DOI:** 10.1101/2025.08.27.672741

**Authors:** Ayame Shimbo, Soichiro Nishiyama, Akane Kusumi, Takuya Morimoto, Hisayo Yamane, Akihiro Itai, Ryutaro Tao

## Abstract

Fruit growth has long been described using single- or double-sigmoid curves; however, these temporal models cannot fully capture the spatial heterogeneity that ultimately shapes a fruit. Here, we present a three-dimensional analysis pipeline that non-destructively tracks spatial fruit growth dynamics from field-collected imaginary. Surface landmarks were drawn, and video recordings were taken throughout development for three pome fruits, apple (*Malus* × *domestica*), Japanese pear (*Pyrus pyrifolia*) and European pear (*Pyrus communis*), and two drupe fruits, peach (*Prunus persica*) and Japanese apricot (*Prunus mume*), to track their motion. Using 3D Gaussian Splatting, we successfully reconstructed 3D models of the fruits, and the landmark displacement could be measured with high accuracy, with R^2^ ≥ 0.98 when compared to manual recordings. We found a common spatial growth gradient in the longitudinal growth shared in the pomes and drupes of the Rosaceae; proximal (stem-end) regions exhibited more pronounced growth than the distal (stylar) end. An exception was found in European pear ‘Bartlett,’ which showed relatively vigorous growth in the distal region, explaining its distinct shape with expanded distal end. Transverse expansion varied far less than longitudinal expansion, with a possible association with initial fruit morphology. Inter-fruit growth variability peaked in the fastest-growing regions, particularly in the distal area of the European pear, highlighting the link between growth vigor and phenotypic variance. These results provide foundational insights into the developmental dynamics of both pome and drupe fruits of the Rosaceae family, contributing to the optimization of fruit size, shape, and uniformity.

## Introduction

Fruit size and shape are key horticultural traits that determine the market value, consumer preference, and post-harvest handling efficiency (Costa et al., 2011; Li et al., 2023). Because agricultural products naturally vary in morphology, products that do not meet the visual standards for these traits are often considered defective (Costa et al., 2011). Therefore, shape uniformity is a critical quality attribute that is ultimately determined by variations in fruit development.

In the Rosaceae family, there are over 100 genera and approximately 3,000 species, including economically important fruits, nuts, ornamental, and timber crops (Shulaev et al., 2008). Rosaceae exhibits diverse fruit types, including drupes, pomes, drupetums, achenes, and achenetums, making it a favored plant family for comparative developmental and evolutionary studies of fruit (Liu et al., 2020). Drupes have fruit flesh derived from the ovary wall and include many agronomically important fruit crops, such as peach (*Prunus persica*), Japanese apricot (*P. mume*), sweet cherry (*P. avium*), and European plum (*P*. x *domestica*). They are also called stone fruits because their endocarp becomes lignified and forms a hard shell that encases one or more seeds. In contrast, pomes, such as apple (*Malus* x *domestica*) and pear (*Pyrus* spp.), have fruit flesh derived primarily from the hypanthium (Pratt, 1988).

Fruit growth is typically modeled as a time-dependent process in one of two major types: a single-sigmoid growth curve, common in pomes (Crane, 1964; Tukey and Young, 1942), and a double-sigmoid growth curve, as observed in drupes (DeJong and Goudriaan, 1989; Nakanishi et al., 1983; Tukey and Young, 1939). The single-sigmoid growth curve is characterized by slow growth at early stages, followed by continuous acceleration during mid-development and gradual deceleration as the fruit reaches maturity. In contrast, the double-sigmoid growth curve is characterized by a lag period after the initial rapid expansion phase, followed by a second growth phase, and a final slowdown as the fruit approaches maturity. In peach, a temporary plateau in fruit growth coincides with endocarp hardening, which is likely caused by the high energy demand required for endocarp lignification (Dardick et al., 2010). Although growth curves can describe the overall growth of an organ well, they are not designed to resolve the localized developmental variations inherent in fruit, which is better characterized through 3D analysis.

Recent advancements in spatially resolved analyses of fruit growth have been particularly notable in model plants. For example, a developmental model has been constructed to explain the morphological differences between *Arabidopsis thaliana* (cylindrical fruit) and *Capsella rubella* (heart-shaped fruit) based on analyses of cell growth rates and division orientations (Eldridge et al., 2016). In contrast, such detailed spatial analyses are still limited in fruit trees. This limitation arises due to the large plant size and extended fruit development period of fruit crops, necessitating specialized measurement techniques and introducing variability, while also presenting challenges in applying molecular genetic approaches similar to those conducted in model species. Consequently, a comprehensive understanding of spatial developmental dynamics in fruit trees lags behind that of model plants.

Fruit shape and development can be quantified using various image-based methods. For example, the cross-sectional shapes of fruits were objectively and efficiently quantified in different species using predefined descriptors or elliptical Fourier descriptors, which are derived from fruit contours extracted from 2D images (Brewer et al., 2006; Currie et al., 2000; Imai et al., 2024; Wang et al., 2024). However, 2D-based methods often fail to represent the full range of shape features because, in most cases, the 2D imaging method acquires shape features that can be seen from only a single perspective of the fruit, which potentially omits biologically important shape features that the original 3D structure possesses. A typical case is the characterization of complex shape development in persimmon (*Diospyros kaki*); complex shape traits such as horizontal and vertical grooves cannot be reliably assessed using 2D methods because of occlusion and covariance with other shape features (Kusumi et al., 2022; Maeda et al., 2018). However, the advent of high-resolution 3D phenotyping pipeline now allows these traits to be captured with precision (Kusumi et al., 2024). Nevertheless, this method still requires imaging in a controlled environment and thus destructive sampling; therefore, continuous, non-destructive 3D measurement of fruit development on the same individual fruit has not been achieved, which ultimately limits the accuracy of spatially resolved growth modeling of the fruit (Kusumi et al., 2025).

Recently, several new paradigms have been established for modeling 3D scenes using machine learning approaches. Neural Radiance Fields (NeRF) is a deep learning-based method for reconstructing 3D scenes by modeling a continuous radiance field that maps each 3D point and viewing direction to color and density, enabling the rendering (synthesis) of images from novel viewpoints (Mildenhall et al., 2021). In recent years, a growing number of studies have applied NeRF to fruit tree research, as it enables high-fidelity reconstruction, even under complex lighting conditions and occlusions (Hu et al., 2024; Peng et al., 2025; Wang et al., 2025). In contrast, 3D Gaussian splatting (3DGS) is an explicit, point-based alternative to NeRF (Kerbl et al., 2023). The major difference between NeRF and 3DGS is their representation: NeRF computes the density and color at each point within the scene, whereas 3DGS represents the scene as a collection of spatially distributed Gaussian functions. As a result, 3DGS can construct 3D models more quickly and with a similar or even higher quality than NeRF. Compared to NeRF, the use of 3DGS in plant phenotyping research remains limited but has recently gained attention (Chen et al., 2025; Shen et al., 2025; Stuart et al., 2025). As both methods can generate high-quality 3D scene models, even from images acquired under outdoor field conditions, they are expected to provide an effective basis for quantitative, full 3D analyses of fruit growth.

In this study, we leveraged machine learning based 3D reconstruction frameworks to continuously and non-destructively track fruit development in the field and to characterize spatial growth heterogeneity in the drupe and pome fruits of Rosaceae. Landmarks were drawn on the fruit surface and their motion on the fruit was recorded on video throughout the growing season. A 3D model of each fruit was then reconstructed, and growth was quantified by tracking the marks over time across different portions of the fruit. Comparable methods have been applied to apple (Skene, 1966), European plum (Khanal et al., 2023) and persimmon (Fujimura, 1935; Kusumi et al., 2025), where they showed vigorous growth in the proximal fruit region. Here, we extended this approach to a broader set of drupes and pomes by incorporating 3D phenotyping. Our approach elucidated the spatial patterns underlying the developmental dynamics shared by drupes and pomes, and provides a fundamental insight and useful pipelines for future research on fruit growth and development.

## Materials and methods

### Plant materials

This study examined three pome species, apple (‘Fuji’), Japanese pear (*Pyrus pyrifolia* ‘Osa Gold’), and European pear (*Pyrus communis* ‘Bartlett’), planted at the Experimental Farm of Kyoto Prefectural University (Seika, Japan), and two drupe species, Japanese apricot ‘Nankou’ and peach ‘Akatsuki’, planted at the Kyoto Farmstead of the Experimental Farm of Kyoto University (Kyoto, Japan). All were managed according to standard horticultural practices. For apple and pear, fruit thinning was carried out until 30–40 days after full bloom (DAFB), leaving one fruit for 4–5 apical buds. In peach, thinning of flower and fruit was carried out accordingly; crop load was limited to 0–1 fruit per short fruiting spur and 1–2 fruits per medium- to long-fruiting shoot by the final thinning at 70 DAFB. No crop load control was applied to Japanese apricot.

### Fruit surface landmarking and field data collection

Landmarks were drawn on the fruit surface using marker ink. The arrangement of the surface landmarking is summarized in Fig 1A. We applied different marking schemes to the pome and drupe. Pome fruits were marked along four stem-stylar lines drawn at 90° intervals. Unlike pome fruits, drupe fruits possess a suture line that introduces additional axis absent in pomes; therefore, they were marked along five stem-stylar lines: two lines on both sides of the suture line, one on the side opposite the suture line, and two between the opposite line and each suture-adjacent line. Three landmarks for pome fruits and four landmarks for drupe fruits were drawn along each line. When the gap between landmarks widened during development, additional landmarks were drawn in between the existing landmarks. In addition, two arbitrarily selected landmarks were drawn in red and manually measured at each observation to verify the measurement accuracy. Fruit growth was recorded once every 1–2 weeks using either an iPhone XR or iPhone 13 (Apple Inc., USA), starting from the stage when the fruit had grown large enough to be marked and continuing until growth ceased. Most of the data collection was conducted during the 2024 growing season. For peach, due to severe fruit drop and insufficient data in 2024, data from both 2024 and 2025 were used for analysis. Each video recording lasted 1–2 min for each fruit, and the fruit was captured from all angles. A ruler was included in each video recording to allow scale calibration. When the tracked fruit dropped, new fruits were selected and marked following the same procedure.

**Figure 1.**
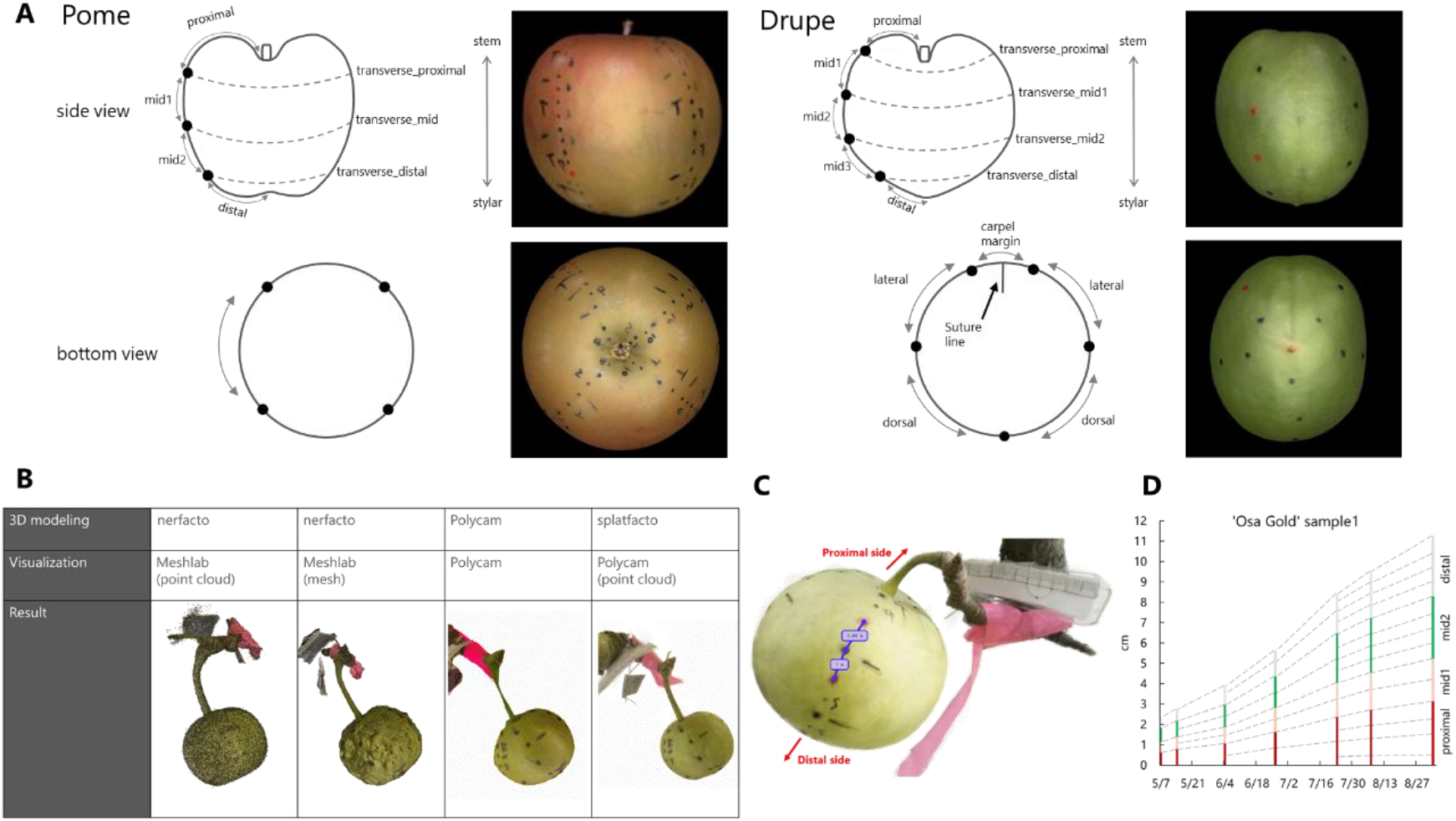
Overview of landmarking methods and 3D modeling for fruit growth analysis. (A) Schematic diagrams of surface landmarking for fruit development analysis. Top: side view, bottom: bottom view. Left: pome, right: drupe. (B) Various combinations of 3D modeling and visualization tools were tested. The best result was obtained using splatfacto from nerfstudio for 3D modeling and Polycam for visualization. (C) Example of distance measurement between surface markers on a fruit using Polycam. After scale calibration with the ruler, the distances between the landmarks were measured. (D) An example of stacked bar graph representing growth in the proximal, mid1, mid2, and distal regions in an ‘Osa Gold’ fruit.

### 3D modeling and phenotyping

Each video recording was trimmed to remove unnecessary segments, such as those in which the fruit was absent or swinging owing to wind. Subsequently, image frames were extracted from the video recordings and aligned using either COLMAP v.3.10 (Schönberger and Frahm, 2016) called from Nerfstudio (Tancik et al., 2023; matching method of *vocab_tree* or *exhaustive*) or Metashape v.2.1.3 (Agisoft; default parameters). The alignment software, matching method, and frame extraction interval were optimized for each video recording according to the alignment quality. Subsequently, the 3D model was reconstructed using the Nerfstudio. In Nerfstudio, either the nerfacto pipeline based on NeRF or the splatfacto pipeline based on 3DGS is used. We also tested Polycam application (Polycam Inc., accessed July 12, 2025, https://poly.cam/) for 3D modeling. Meshes generated by nerfacto were visualized using MeshLab 2023.12 (Cignoni et al., 2008), whereas splat data generated by splatfacto were visualized using MeshLab or Polycam. Some video recordings were excluded from the analysis because 3D reconstruction was not possible, likely because of object motion caused by wind or incomplete view coverage.

Scale calibration was performed using the ruler included in the reconstructed 3D models using the scale calibration function of the Polycam. In cases where the ruler was not reconstructed, two red marks whose distances had been measured manually at each recording were used as a reference scale. The distances between the landmarks were then measured using the length measurement function of the Polycam.

### Data analysis

For longitudinal fruit development analysis, the distances between successive landmarks along a single stem-stylar line that included red marks were measured. The distances within each defined region were then summed to calculate regional length. For pomes, the region from the fruit-pedicel junction to the first landmark was designated as the proximal region, the region between the first and second landmarks as mid1, between the second and third landmarks as mid2, and between the third landmark and the fruit apex as the distal region (Fig 1A). Drupes were partitioned in the same manner, but with one extra landmark, creating an additional mid3 region between mid2 and the distal region (Fig 1A). The regional lengths were then used to calculate the growth rate, which was expressed as a fold-change per week, calculated as Fold-change per week 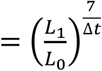,

where *L*_0_ is the length of the previous observation, *L*_1_ is the length of the current observation, and Δ*t* is the number of days between the two measurements. Measurement dates for sample are listed in Table S1, which also flags missing observations caused by physiological fruit drop or sub-optimal reconstruction quality under certain imaging conditions.

For the transverse growth analysis, only the original landmarks (not additional landmarks) were considered. For pomes, three landmarks, (transverse_proximal, transverse_mid, and transverse_distal), were marked on each of the four stem-stylar lines, yielding four measurements per position (Fig 1A); their average was used for further analysis. For drupes, five stem-style lines, for each with four landmarks (transverse_proximal, transverse_mid1, transverse_mid2, and transverse_distal), were grouped into three anatomical portions: carpel margin (along the suture line), lateral, and dorsal (opposite the suture line) sides (see Fig 1A for a diagram). Two values were obtained for each of the lateral and dorsal regions, and their averages were used for the analysis. If a 3D model was incomplete and lacked full 360° coverage, values from the available fruit portion were retained. The fold-change per week was calculated as described for the longitudinal growth analysis.

Subsequently, an allometric analysis was conducted to evaluate the relative growth dynamics of each fruit region in relation to total fruit length (Huxley, 1932; Skene, 1966). This analysis was restricted to marked fruits that remained on the tree and were fully imaged throughout the entire growing season. Regional and total lengths were log_10_-transformed, and for each sample the relationship

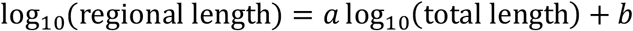

was fitted by linear regression, where *a* is the slope, and *b* is the intercept. This single linear model adequately described the allometric relationship between regional and total fruit lengths in most species; however, the distal region of apple and peach deviated from linearity (see Results). To accommodate this, piecewise linear regression was applied. For each candidate breakpoint *c* (expressed as log_10_ total length), the data were split into two subsets at *c*, and a piecewise linear model was fitted to each segment as:

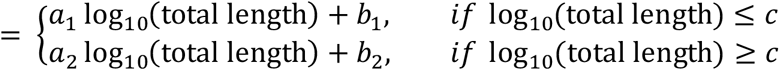

The breakpoint *c* was iteratively shifted across the range of total lengths, and the one yielding the highest mean R^2^ across the two segments was selected.

To quantify the inter-fruit variation, each distance value was normalized by setting the landmark distance on the latest common start date to 1. As the measurement dates differed among the samples, the interval between successive observations was uneven. Therefore, spline interpolation was applied to estimate the daily landmark distances. The standard deviation among the fruits over this relative distance was then calculated for each day until the earliest common end date.

## Results

### Evaluation of in-field 3D fruit measurement methods

The quality of the visualization was evaluated using different combinations of 3D modeling and rendering methods. Among them, the combination of 3D modeling with splatfacto and visualization with Polycam enabled a faster workflow of visually high-quality models (Fig 1B). Using this method, the developmental trajectories of the five fruit species were tracked non-destructively (Supplementary Fig S1). The temporal changes in fruit morphology were successfully captured throughout the developmental timeline. Thereafter, growth in each region was confirmed to be tracked continuously using surface landmarks throughout development in the same sample (Fig 1C and D). Concordance with the manual caliper measurements and with 3DGS-based measurements was high for all species, with coefficients of determination (R^2^) ≥ 0.98 (Supplementary Fig S2).

Using this method, the longitudinal growth rates among the five species were compared (Fig 2). In all species, growth was vigorous during the early fruit development stage, followed by a gradual slowdown toward maturity, consistent with typical fruit growth patterns.

**Figure 2.**
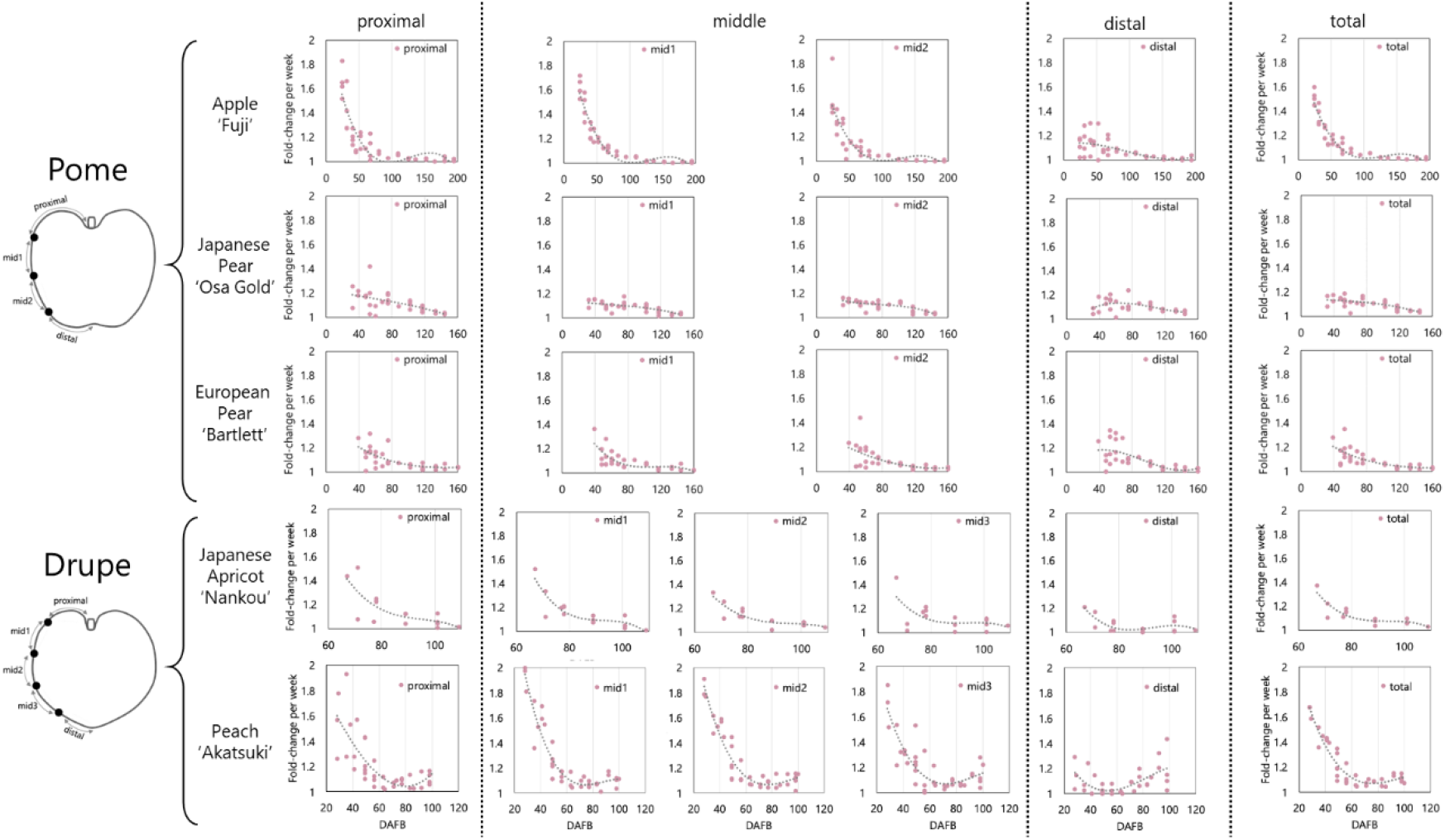
Spatial analysis of longitudinal fruit growth. From left to right: proximal, middle (pome: mid1 and mid2; drupe: mid1, mid2, and mid3), distal, and total (the sum of all regions) values. From top to bottom: Apple ‘Fuji’, Japanese pear ‘Osa Gold’, European pear ‘Bartlett’, Japanese apricot ‘Nankou’, and Peach ‘Akatsuki’.

### Comparative analysis of growth rates across fruit regions in pome

In pome fruits, the temporal pattern of longitudinal growth rates was generally similar and declined from the proximal to the distal region, although the pattern and magnitude varied considerably among the species (Fig 2). In apple ‘Fuji,’ the growth differed markedly between the proximal and distal regions. In particular, growth was more vigorous in the proximal and mid1 regions and slower in the distal region. In Japanese pear ‘Osa Gold,’ although the growth gradient was relatively smaller, growth in the proximal region was most vigorous, consistent with the pattern observed in apple ‘Fuji.’ In contrast, in European pear ‘Bartlett,’ the regional differences in growth were subtle. Compared to apple ‘Fuji’ and Japanese pear ‘Osa Gold,’ relatively vigorous growth was observed in the distal region in European pear ‘Bartlett,’ being consistent with its enlarged appearance of the distal portion and a larger diameter than that of the proximal region (Supplementary Fig S1).

In contrast to longitudinal growth, spatial variation in transverse growth was relatively modest (Fig 3). In apple ‘Fuji,’ longitudinal growth in the distal region was clearly less than that in the proximal, mid1, and mid2 regions, whereas transverse growth in the distal region (transverse_distal) was not as limited as in the longitudinal direction and was moderately vigorous. In Japanese pear ‘Osa Gold’ and European pear ‘Bartlett,’ the growth patterns among regions were similar between longitudinal and transverse directions; In ‘Osa Gold,’ transverse growth was most vigorous in the proximal region (transverse_proximal), as also seen in the longitudinal direction. In ‘Bartlett,’ distal region (transverse_distal) also exhibited vigorous transverse growth.

**Figure 3.**
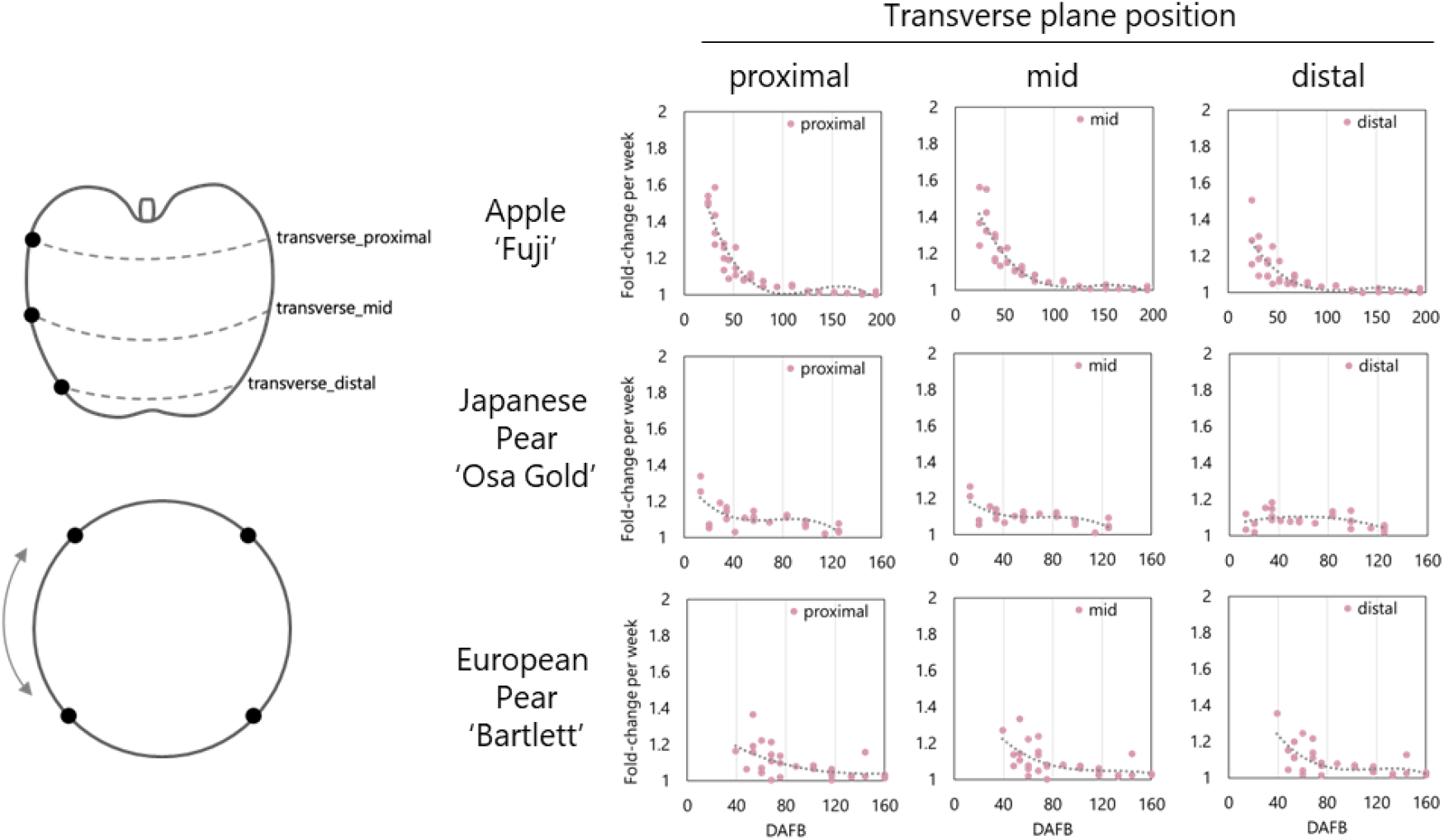
Spatial analysis of transverse growth of pomes. From top to bottom: Apple ‘Fuji’, Japanese pear ‘Osa Gold’, and European pear ‘Bartlett’. From left to right: transverse_proximal, transverse_mid, and transverse_distal regions.

### Comparative analysis of growth rates across fruit regions in drupe

The longitudinal growth patterns were similar between Japanese apricot ‘Nankou’ and peach ‘Akatsuki,’ in which the proximal region showed more vigorous growth than the distal region (Fig 2). However, the temporal patterns of these two species were different. In Japanese apricot ‘Nankou,’ growth plateaued during the late stage of fruit development, whereas in peach ‘Akatsuki,’ growth was stalled at the middle stage but accelerated again at the late stage. Furthermore, in the distal region of peach ‘Akatsuki,’ growth was more vigorous in the late stage than in the early developmental stage.

For transverse growth in drupes, spatial analysis was conducted with a focus on the suture line, which serves as a structural axis of growth that was not present in the pomes (Fig 4). The overall growth pattern resembled that observed in the longitudinal direction, with peach ‘Akatsuki’ exhibiting increased growth during the late developmental stage, whereas Japanese apricot ‘Nankou’ did not. In ‘Nankou,’ in contrast to the longitudinal analysis, the distal region (transverse_distal) exhibited transverse growth comparable to other regions. In peach ‘Akatsuki,’ unlike Japanese apricot ‘Nankou,’ the transverse_distal region showed limited growth compared with the other regions especially during the early developmental stage, consistent with the longitudinal growth trend (Figs 2 and 4). However, during the late developmental stage, recovery of transverse growth was observed particularly in the distal region (transverse_distal). In both species, the regional growth rates along the transverse axis were generally uniform, and no obvious growth variation along the suture line was observed in this study.

**Figure 4.**
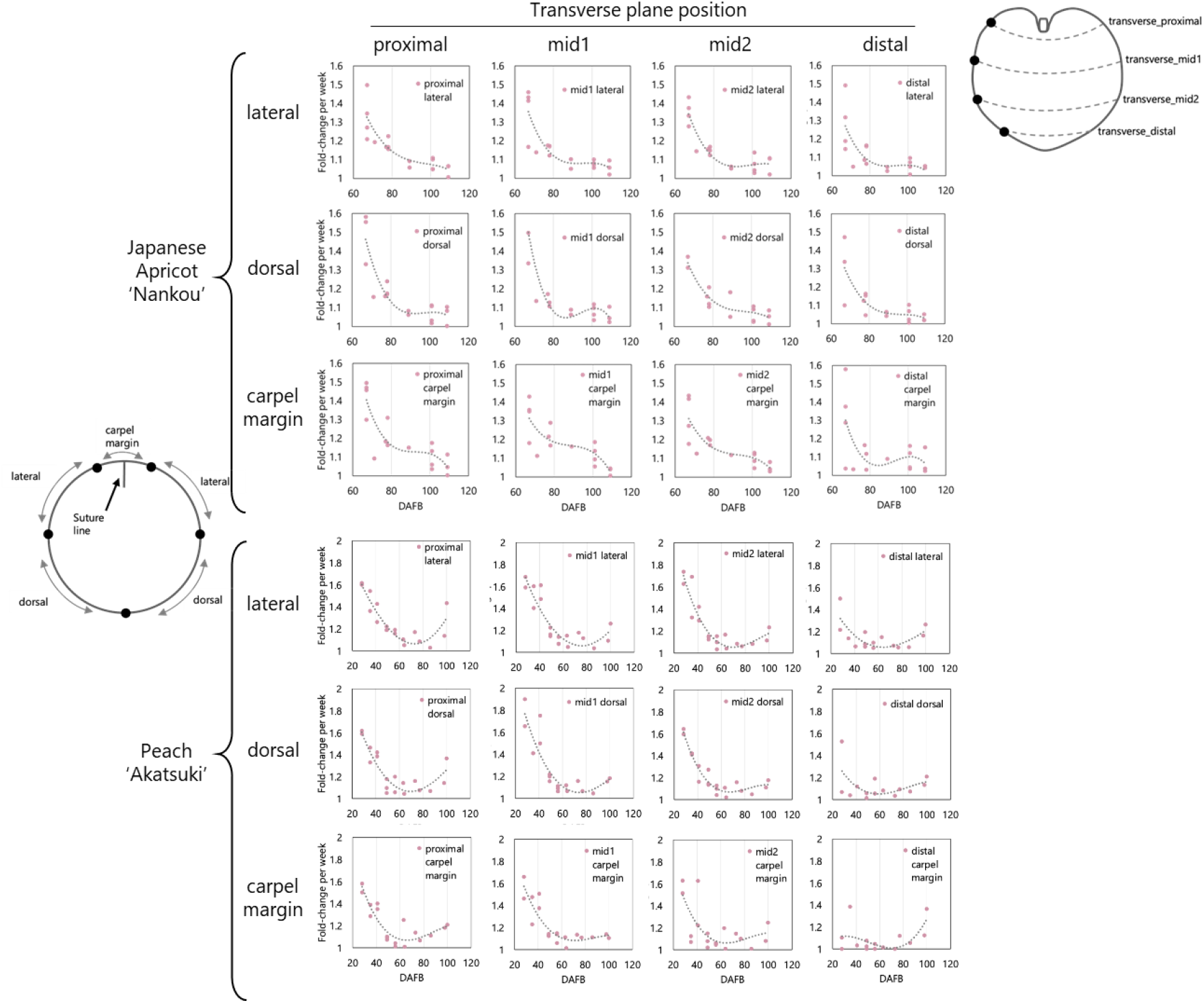
Spatial analysis of transverse growth of drupes. From top to bottom: Japanese apricot ‘Nankou’, and peach ‘Akatsuki’, within each species: lateral, dorsal, and carpel margin regions. From left to right: transverse_proximal, transverse_mid1, transverse_mid2, and transverse_distal regions.

### Allometric analysis of spatially localized fruit growth

Allometric relationships in regional fruit growth have long been suggested (Barabé and Jean, 1995; Skene, 1966); here we observed allometric relationships between regional growth and total fruit length for each species and region (Supplementary Figs. 3–7). Across species, hyperallometric growth (*a* > 1), in which regional growth was higher relative to the overall fruit, was detected in specific regions (Table 1): proximal and mid1 of apple ‘Fuji,’ proximal and distal of Japanese pear ‘Osa Gold,’ distal of European pear ‘Bartlett,’ proximal-mid2 of Japanese apricot ‘Nankou,’ and mid1-mid3 of peach ‘Akatsuki.’ These findings agree with the absolute growth rate analysis (Fig 2), particularly pronounced growth in the distal region of ‘Bartlett.’

**Table 1.**
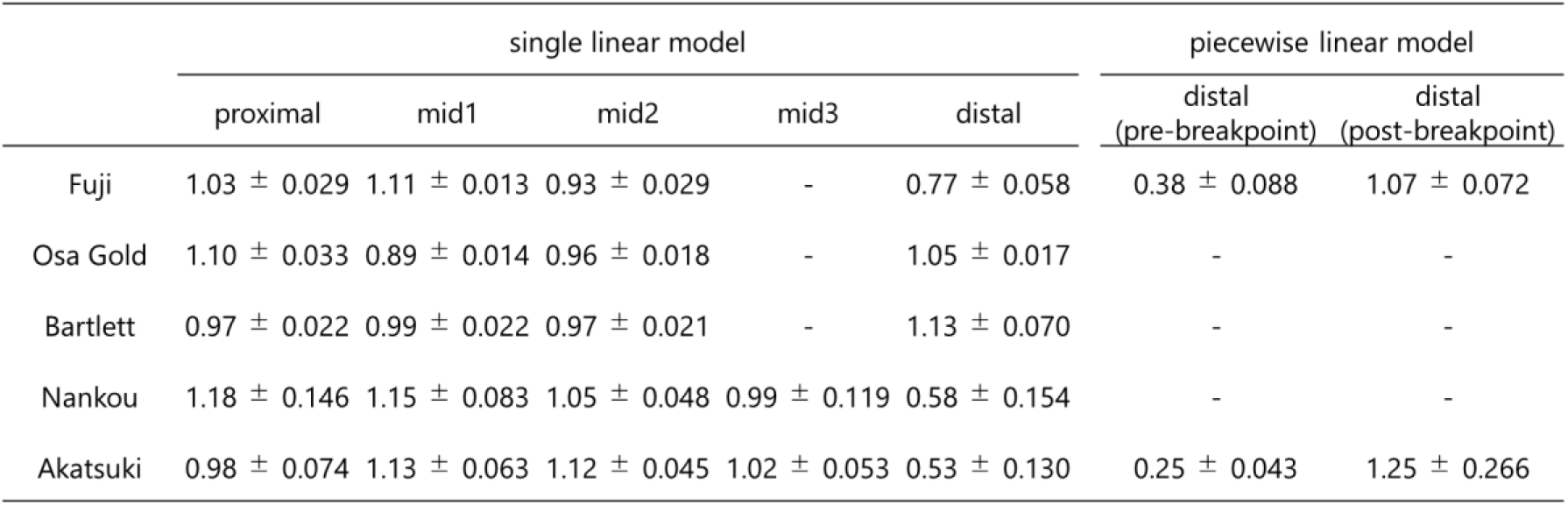
Summary of regional allometry. Regression slopes (*a*) are shown as mean ± SE (*N* = 4-5) for each species and region. For the piecewise linear model, slopes are presented separately as pre-breakpoint (≤ *c*) and post-breakpoint (≥ *c*).

Linear regression fittings were satisfactory for most species and regions, but distal regions of apple ‘Fuji’ and peach ‘Akatsuki’ showed poor fits (R^2^=0.66 at the lowest; Supplementary Figs 3A and 7A), suggesting a slope shift during development. Piecewise regression (Supplementary Figs 3B and 7B) improved the fitness and revealed a constant trend: an early hypoallometric phase (*a* < 1), in which slower regional growth relative to total fruit length, followed by a late hyperallometric phase (*a* > 1). The breakpoint occurred at the total length of 5-8 cm for ‘Fuji’ and around 8 cm for ‘Akatsuki,’ marking a transition in distal growth dynamics. Collectively, these results identify species- (or cultivar-) specific “growth-hotspots” and a critical developmental switch from slower- to faster-than-proportional-expansion in the distal region.

### Spatial analysis of inter-fruit growth variations

Inter-fruit variations in growth trajectories have substantial economic implications, since they influence the fruit uniformity. To quantify these differences at spatial resolution, we applied spline interpolation to estimate daily growth and then normalized each trajectory (see Materials and Methods). The resulting smooth curves enabled day-by-day comparisons of growth among fruits (Supplementary Figs S8-12). The regions showing higher growth variation differed by species (Fig 5). In apple ‘Fuji,’ Japanese pear ‘Osa Gold,’ and Japanese apricot ‘Nankou,’ the variation was greatest in the proximal region. In peach ‘Akatsuki,’ the variation was greatest in both the proximal and mid1 regions. In contrast, European pear ‘Bartlett’ showed the greatest variation in the distal region. These regions matched with the regions that exhibited vigorous growth (Fig 2), implying that faster growing regions are intrinsically more prone to inter-fruit variation.

**Figure 5.**
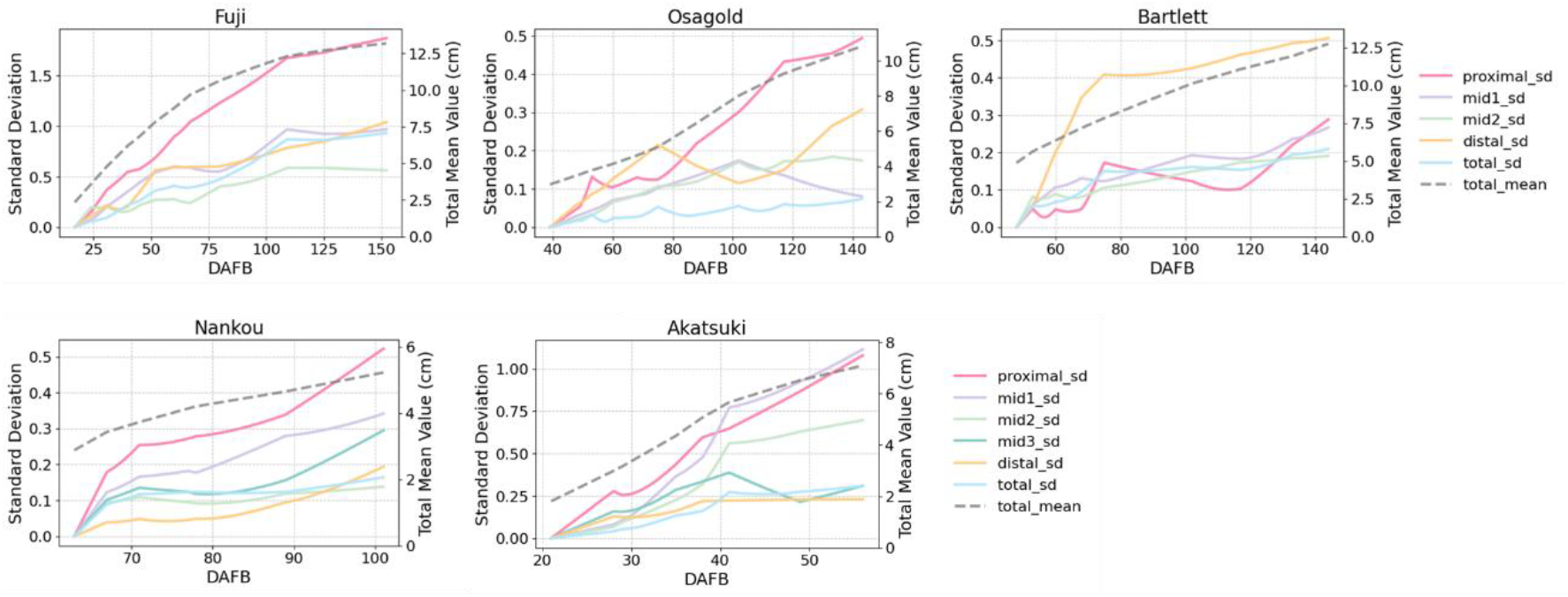
Inter-fruit spatial variation in growth. (A) Pomes (from left to right: apple ‘Fuji’, Japanese pear ‘Osa Gold’, and European pear ‘Bartlett’). (B) Drupe (left: Japanese apricot ‘Nankou’; right: peach ‘Akatsuki’). The dashed line indicates the total mean length (cm), and the solid lines indicate the standard deviation for each region.

## Discussion

We observed that fruit growth was more vigorous in the proximal region and slower in the distal region in four of the five species (cultivars) examined in this study (Fig 2 and Table 1). This proximal-dominant pattern appeared in both pomes and drupes and aligns with previous reports on the European plum, which also documented enhanced growth near the stem end (Khanal et al., 2023). A classic study in apple showed maximum growth in the middle region and the least growth in the distal region (Skene, 1966), but it divided the fruit surface into six regions, whereas our scheme used four. Despite the difference in regional partitions, the overall conclusions are comparable. A similar gradient has been reported in persimmon, a species phylogenetically distant from Rosaceae (Fujimura, 1935; Kusumi et al., 2025). Future studies applying similar spatial 3D phenotyping apprioach to a wider range of species may help to reveal whether the proximal-dominant growth is conserved across fleshy fruits. In parallel, intraspecific variation in fruit morphology has been well characterized in 2D in various species, including plum (*P. salicina*) (Xia et al., 2023), peach (Guo et al., 2018), apple (Dar et al., 2015) and melon (*Cucumis melo*) (Stepansky et al., 1999); expanding such analyses and resources into the spatial 3D growth are warranted, as it may help reveal the developmental mechanisms driving shape variation.

Among the five species used in this study, European pear ‘Bartlett’ exhibited a unique growth pattern in which the distal region showed relatively vigorous growth (Fig 2). In addition, the variation in growth was greatest in the distal region, unlike in other species (Fig 5). These growth features may be related to the characteristic fruit shape of ‘Bartlett.’ Because the fruit shape of ‘Bartlett’ changes considerably throughout development (Bain, 1961) and tends to lack axial symmetry (Supplementary Fig S1), 2D-based metrics, including second derivative-based quantification of pyriform features

(White et al., 2000) and elliptical Fourier descriptors of the fruit section (Wang et al., 2024), provide only a partial description of these morphological changes. Our pipeline complements these approaches by capturing more complex and dynamic morphological characteristics. Given the wide range of shape variations in European pears, including globose, ovate, conical, cylindrical, spindle-shaped, pear-shaped, and inverted heart-shaped forms driven by genotype, nutrition, and environmental influences (Wang et al., 2024; Bayazit et al., 2016), precise and dynamic morphometric methodologies, as exemplified in the present study, may provide valuable insights into the mechanisms underlying shape diversity.

Despite the similarity in fruit shapes, apple ‘Fuji’ and Japanese pear ‘Osa Gold’ showed different growth patterns especially in the transverse axis (Figs 2 and 3). This difference was likely attributable to variations in fruit morphology during early stages. ‘Fuji’ exhibited an elongated shape at the early developmental stage, which can be subsequently modified by pronounced transverse growth, culminating in a more rounded form (Supplementary Fig S1). In contrast, ‘Osa Gold’ possessed a naturally rounded morphology from the beginning of development and was considered to exhibit comparatively limited transverse expansion (Supplementary Fig S1). Ultimately, the fruits of both species are likely to have a rounded morphology.

In the allometric analysis, we observed growth breakpoints that result in more active growth in the later stage within the distal region of apple ‘Fuji’ and peach ‘Akatsuki’ (Table 1). This pattern may suggest that, in the distal part, cell division is not particularly active during the early developmental stages, whereas cell expansion occurs at a similar or even greater level than in other regions during the later stages. One possible explanation is that signals promoting cell division may originate from the proximal (stem-end) side. Although this remains speculative, these findings highlight a potential developmental mechanism that warrants further experimental validation.

The method developed in this study enabled continuous and non-destructive tracing of fruit growth throughout development and is expected to contribute to elucidating the mechanisms underlying physiological disorders caused by fruit growth gradients, such as cracking and shriveling. For instance, in sweet cherry (*P. avium*), larger fruits tend to crack more easily, and kidney- or heart-shaped varieties with deeper stem cavities retain water longer, thereby increasing the risk of cracking (Simon, 2006). In apple, cultivars exhibiting more vigorous transverse growth during fruit enlargement tend to be more prone to fruit cracking (Yamamoto et al., 1993). In European plum, late-stage rapid growth in the proximal region caused neck shriveling (Khanal et al., 2023). By visualizing spatio-temporal growth, our approach may be applied to dissect such relationships to minimize physiological disorders in the future.

The method developed in this study has three main limitations. First, it could not capture the earliest developmental stages of fruit growth because physical landmarks can only be drawn once the fruit reaches a certain size. Second, although early-stage fruits could still be reconstructed using detectable landmarks (Supplementary Fig S1), the resolution was limited, likely due to the difficulty of reconstructing small objects with 3DGS. Third, in orchard environments, occlusions by branches, leaves, and fruits led to insufficient image data, resulting in difficulties in recovering local details and increased aliasing, as previously noted (Chen et al., 2025). To overcome these limitations, further improvements in the accuracy and robustness of imaging method are required. Moreover, by developing this method into a predictive growth model, it may be possible to visualize spatial variations in growth, without relying on physical landmarks. Such advancements are expected to enable more accurate analyses, even for small fruits.

## Conclusion

The 3DGS-based method employed in this study enables the reconstruction of high-quality 3D models of fruit that are both visually realistic and quantitatively accurate, as validated against caliper measurements. This approach allows continuous, nondestructive, region-specific tracking of growth at high precision, including the precise localization of surface features. In Rosaceae, growth is generally more vigorous on the proximal side and slower on the distal side. However, in the European pear, the distal region exhibited relatively vigorous growth, which may have contributed to its unique shape. In addition, the greatest variation in growth among the samples was observed in regions where growth was the most vigorous. These findings may contribute to a better understanding of spatially resolved growth dynamics and morphological diversity of fruit across horticulturally important crops.

## Acknowledgment

This research was supported by the Japan Society for the Promotion of Science KAKENHI (grant no. 21KK0269 to SN and no. 24KJ1497 to AK), the JST BOOST Program (grant no. JPMJBY24F7 to SN) and a research grant from the Hirose Foundation to SN.

## Author Contributions

SN conceived and designed the study. AS and AK optimized field data collection and collected the data. AS and SN developed the methodology, analyzed the data, and drafted the manuscript. SN, TM, HY, AI and RT prepared plant materials and experimental facilities. All authors read and approved the final version of the manuscript.

