## Supplementary Materials for "Spatially resolved analysis of growth dynamics in pome and drupe fruits of Rosaceae using 3D Gaussian Splatting"

Table S1. Overview of samples, measurement dates, and analyses applied.

| | | Analysis | | | | Measurement Dates | | | | | | | | | | |
|---|---|---|---|---|---|---|---|---|---|---|---|---|---|---|---|---|
| | | Longitudinal growth | Transverse growth | Allometry | Variation | | | | | | | | | | | |
| Fuji | sample1 | ✓ | ✓ | ✓ | ✓ | 2024/5/7 | 2024/5/14 | 2024/5/21 | 2024/5/30 | 2024/6/19 | 2024/6/26 | 2024/7/23 | 2024/9/3 | 2024/9/19 | 2024/10/17 | |
| | sample2 | ✓ | ✓ | ✓ | ✓ | 2024/5/7 | 2024/5/14 | 2024/5/30 | 2024/6/4 | 2024/6/11 | 2024/6/19 | 2024/6/26 | 2024/7/9 | 2024/8/7 | 2024/10/31 | |
| | sample3 | ✓ | ✓ | ✓ | ✓ | 2024/5/7 | 2024/5/14 | 2024/5/21 | 2024/5/30 | 2024/6/11 | 2024/7/9 | 2024/8/23 | 2024/10/2 | 2024/10/17 | 2024/10/31 | |
| | sample5 | ✓ | ✓ | ✓ | ✓ | 2024/5/7 | 2024/5/21 | 2024/5/30 | 2024/6/4 | 2024/6/26 | 2024/8/7 | 2024/9/19 | | | | |
| | sample7 | ✓ | ✓ | ✓ | ✓ | 2024/5/7 | 2024/5/14 | 2024/5/21 | 2024/6/11 | 2024/6/26 | 2024/8/23 | 2024/10/17 | | | | |
| Osa Gold | sample1 | ✓ | ✓ | ✓ | ✓ | 2024/5/21 | 2024/6/26 | 2024/7/23 | 2024/8/7 | 2024/8/23 | 2024/9/3 | | | | | |
| | sample2 | ✓ | ✓ | ✓ | ✓ | 2024/5/14 | 2024/5/30 | 2024/6/4 | 2024/6/11 | 2024/6/26 | 2024/7/23 | 2024/8/7 | 2024/9/3 | | | |
| | sample3 | ✓ | ✓ | ✓ | ✓ | 2024/5/14 | 2024/6/4 | 2024/6/26 | 2024/7/23 | 2024/8/7 | 2024/9/3 | | | | | |
| | sample5 | ✓ | ✓ | ✓ | ✓ | 2024/5/7 | 2024/5/14 | 2024/5/21 | 2024/5/30 | 2024/6/4 | 2024/6/11 | 2024/6/26 | 2024/7/23 | 2024/8/7 | 2024/8/23 | 2024/9/3 |
| | sample7 | ✓ | ✓ | ✓ | ✓ | 2024/5/7 | 2024/5/14 | 2024/6/4 | 2024/6/19 | 2024/6/26 | 2024/7/9 | 2024/7/23 | 2024/8/7 | 2024/9/3 | | |
| Bartlett | sample1 | ✓ | ✓ | ✓ | ✓ | 2024/5/14 | 2024/5/30 | 2024/6/11 | 2024/6/26 | 2024/7/23 | 2024/8/7 | 2024/9/3 | | | | |
| | sample2 | ✓ | ✓ | ✓ | ✓ | 2024/5/7 | 2024/5/21 | 2024/5/30 | 2024/6/4 | 2024/6/11 | 2024/6/26 | 2024/7/9 | 2024/7/23 | 2024/8/7 | 2024/9/3 | 2024/9/19 |
| | sample3 | ✓ | ✓ | ✓ | ✓ | 2024/5/14 | 2024/5/30 | 2024/6/4 | 2024/6/11 | 2024/6/19 | 2024/6/26 | 2024/7/23 | 2024/8/7 | 2024/8/23 | 2024/9/19 | |
| | sample7 | ✓ | ✓ | ✓ | ✓ | 2024/5/30 | 2024/6/4 | 2024/6/11 | 2024/6/19 | 2024/8/7 | 2024/8/23 | 2024/9/3 | 2024/9/19 | | | |
| Nankou | sample2 | ✓ | ✓ | ✓ | ✓ | 2024/4/18 | 2024/4/22 | 2024/4/26 | 2024/5/26 | | | | | | | |
| | sample3 | ✓ | ✓ | ✓ | ✓ | 2024/4/18 | 2024/4/22 | 2024/4/26 | 2024/5/2 | 2024/5/14 | 2024/5/26 | 2024/6/3 | | | | |
| | sample5 | ✓ | ✓ | ✓ | ✓ | 2024/4/18 | 2024/5/3 | 2024/5/14 | 2024/5/26 | | | | | | | |
| | sample6 | ✓ | ✓ | ✓ | ✓ | 2024/4/22 | 2024/5/3 | 2024/5/14 | 2024/5/26 | | | | | | | |
| Akatsuki | sample1 | ✓ | ✓ | ✓ | ✓ | 2024/4/25 | 2024/5/3 | 2024/5/12 | 2024/5/23 | 2024/5/30 | 2024/6/6 | 2024/6/20 | 2024/7/11 | | | |
| | samlpe4 | ✓ | | ✓ | | 2024/4/30 | 2024/5/23 | 2024/6/14 | 2024/6/20 | 2024/6/26 | 2024/7/5 | 2024/7/11 | | | | |
| | sample8 | ✓ | | ✓ | | 2024/5/14 | 2024/5/17 | 2024/5/23 | 2024/6/6 | 2024/6/20 | 2024/7/5 | 2024/7/11 | | | | |
| | sample10 | ✓ | | ✓ | | 2024/5/14 | 2024/5/23 | 2024/5/30 | 2024/6/14 | 2024/6/26 | 2024/7/11 | | | | | |
| | sample2025_5 | ✓ | ✓ | ✓ | ✓ | 2025/4/30 | 2025/5/7 | 2025/5/14 | 2025/5/20 | 2025/5/28 | 2025/6/4 | | | | | |
| | sample2025_10 | ✓ | ✓ | ✓ | ✓ | 2025/4/30 | 2025/5/7 | 2025/5/14 | 2025/5/20 | 2025/5/28 | 2025/6/4 | 2025/6/12 | 2025/6/21 | 2025/7/4 | 2025/7/18 | |

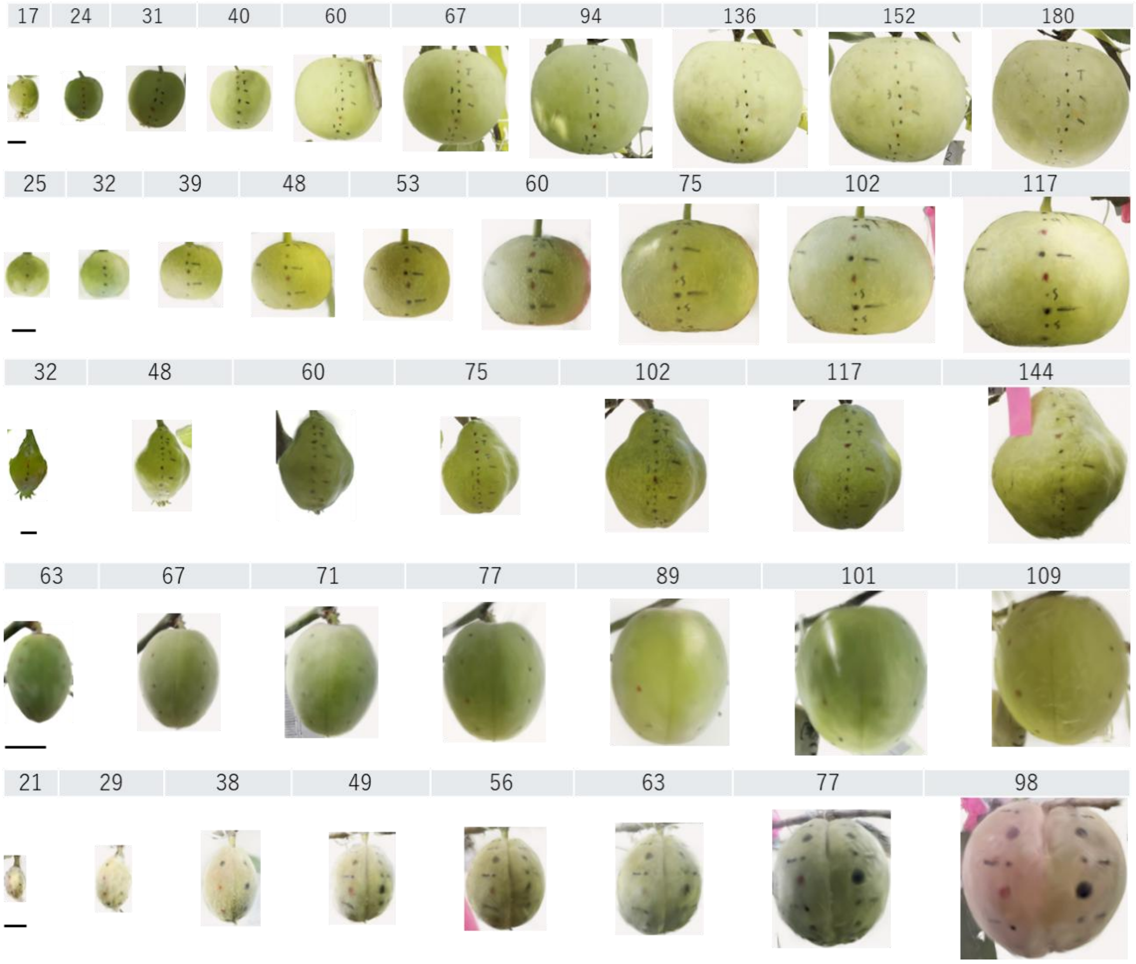


Figure S1. Representative view of 3D models of pome and drupe fruits tracked across development. From top to bottom: Apple ‘Fuji’, Japanese pear ‘Osa Gold’, European pear ‘Bartlett’, Japanese apricot ‘Nanko’ and peach ‘Akatsuki’. The numbers above the images indicate days after full bloom (DAFB). Bar=1cm.

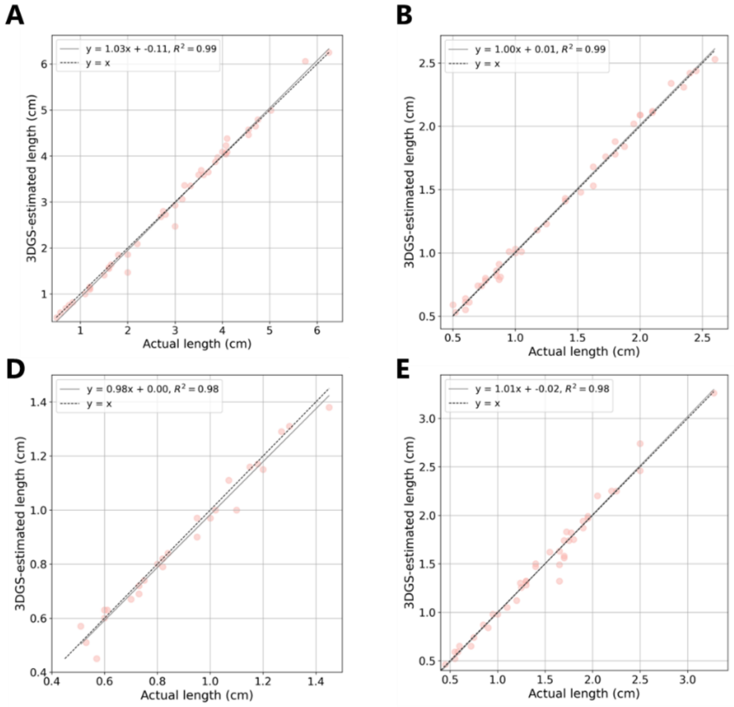


Figure S2. Accuracy of 3D measurements. The actual length was hand-measured using a caliper. A: 'Fuji', B: 'Osa Gold', C: 'Bartlett', D: 'Nankou', E: 'Akatsuki'. The solid line represents the regression line, and the dashed line represents the identity line (y=x).

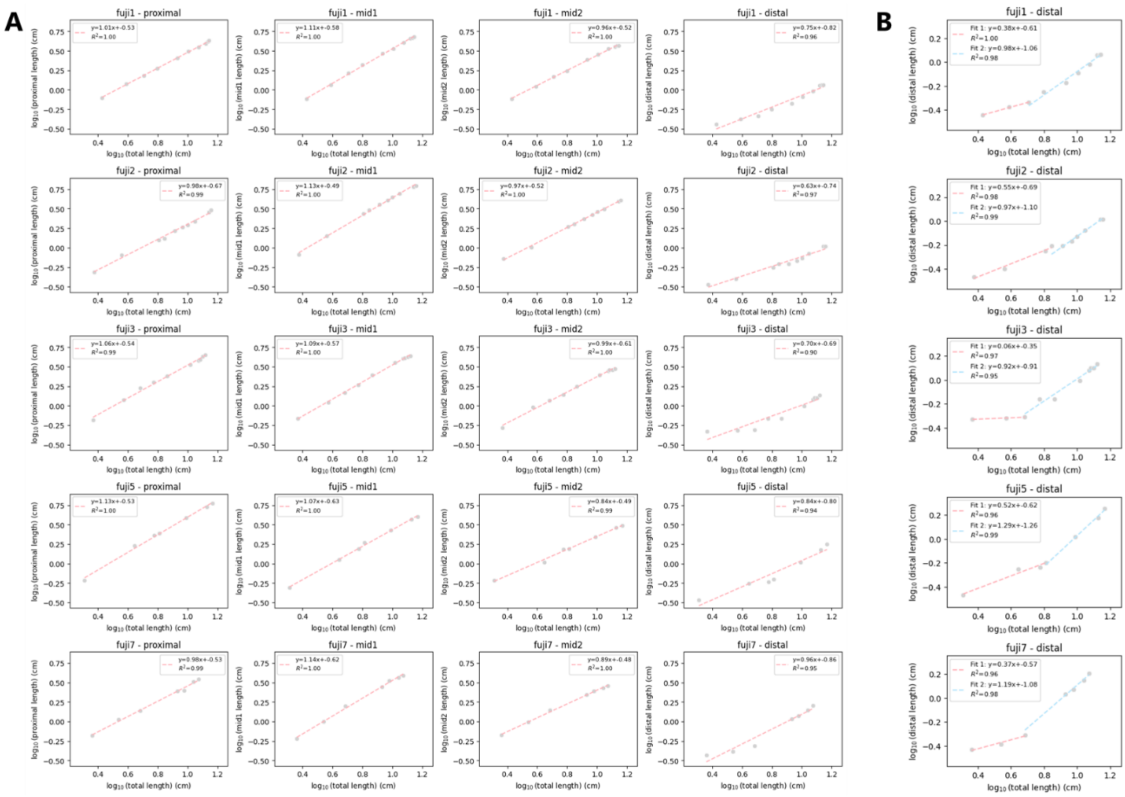


Figure S3. Allometric analysis of spatial growth in apple 'Fuji'. (A) Single linear regression applied to all regions. From left to right: proximal, mid1, mid2, and distal regions. (B) Piecewise linear regression applied only to the distal region.

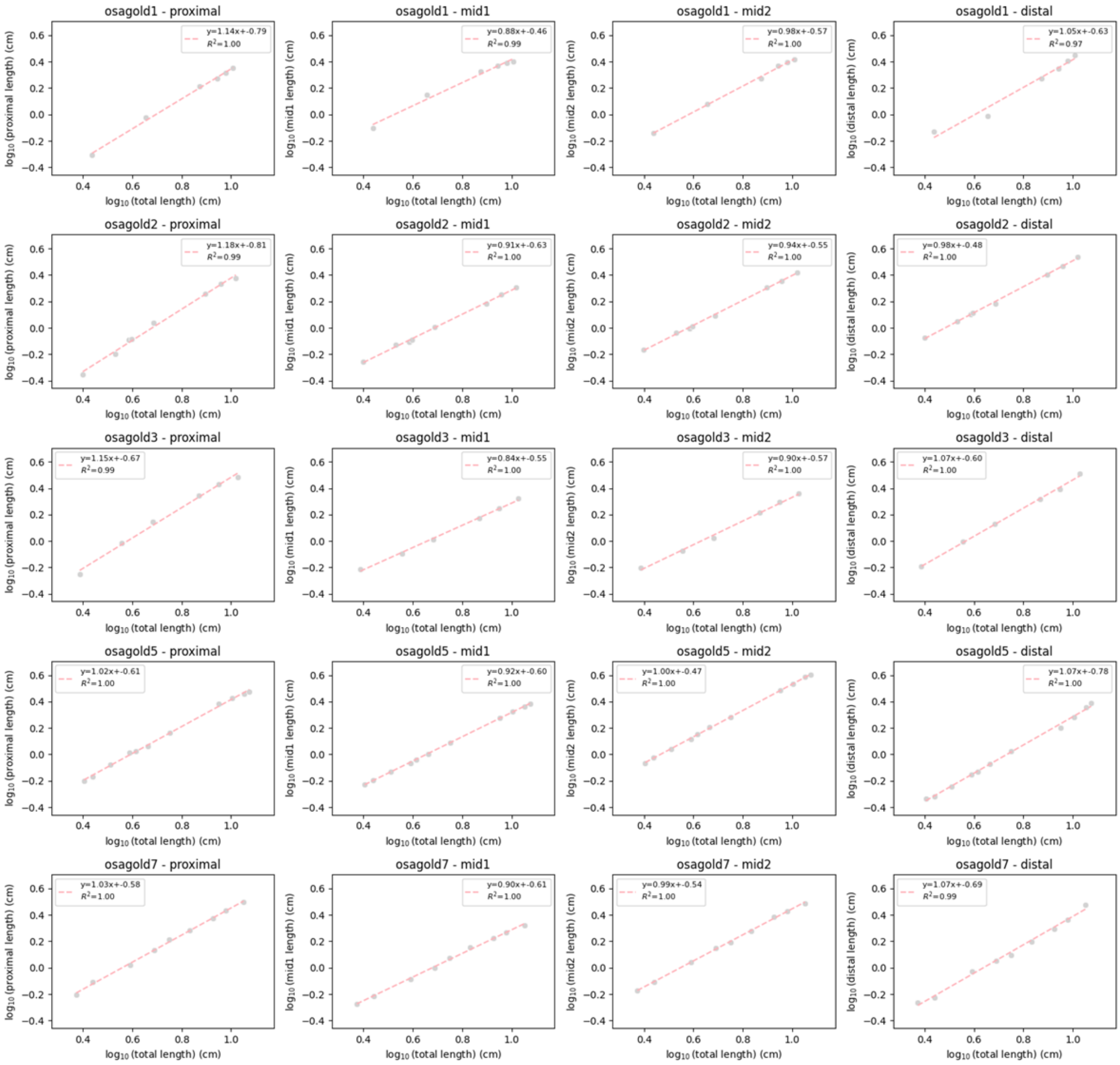


Figure S4. Allometric analysis of spatial growth in Japanese pear ‘Osa Gold’. From left to right: proximal, mid1, mid2, and distal regions.

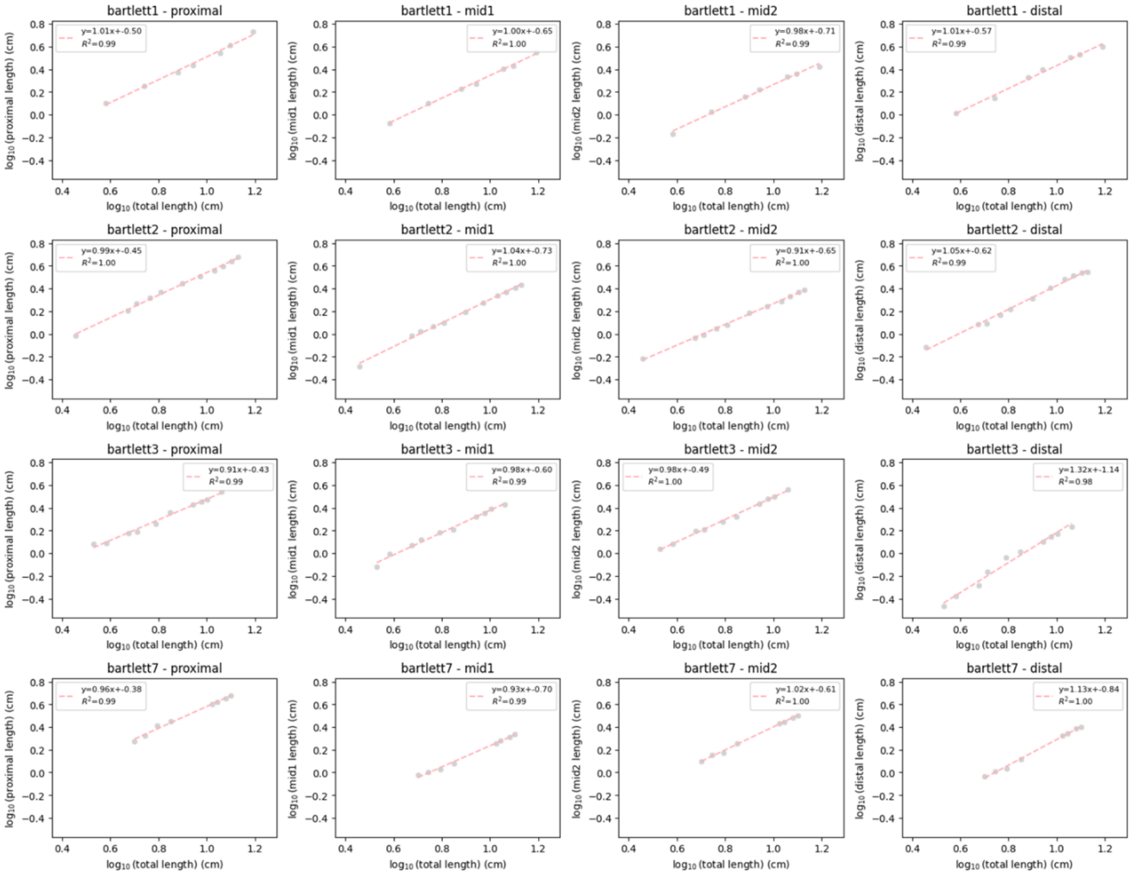


Figure S5. Allometric analysis of spatial growth in European pear ‘Bartlett’. From left to right: proximal, mid1, mid2, and distal regions.

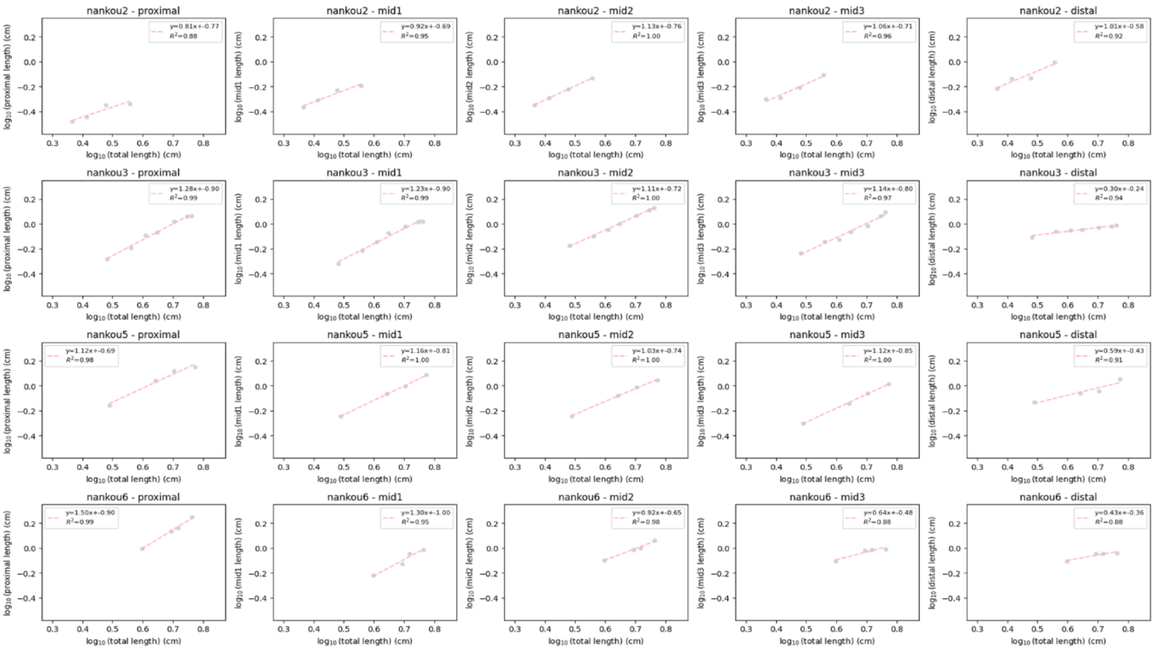


Figure S6. Allometric analysis of spatial growth in Japanese apricot 'Nankou'. From left to right: proximal, mid1, mid2, mid3 and distal regions.

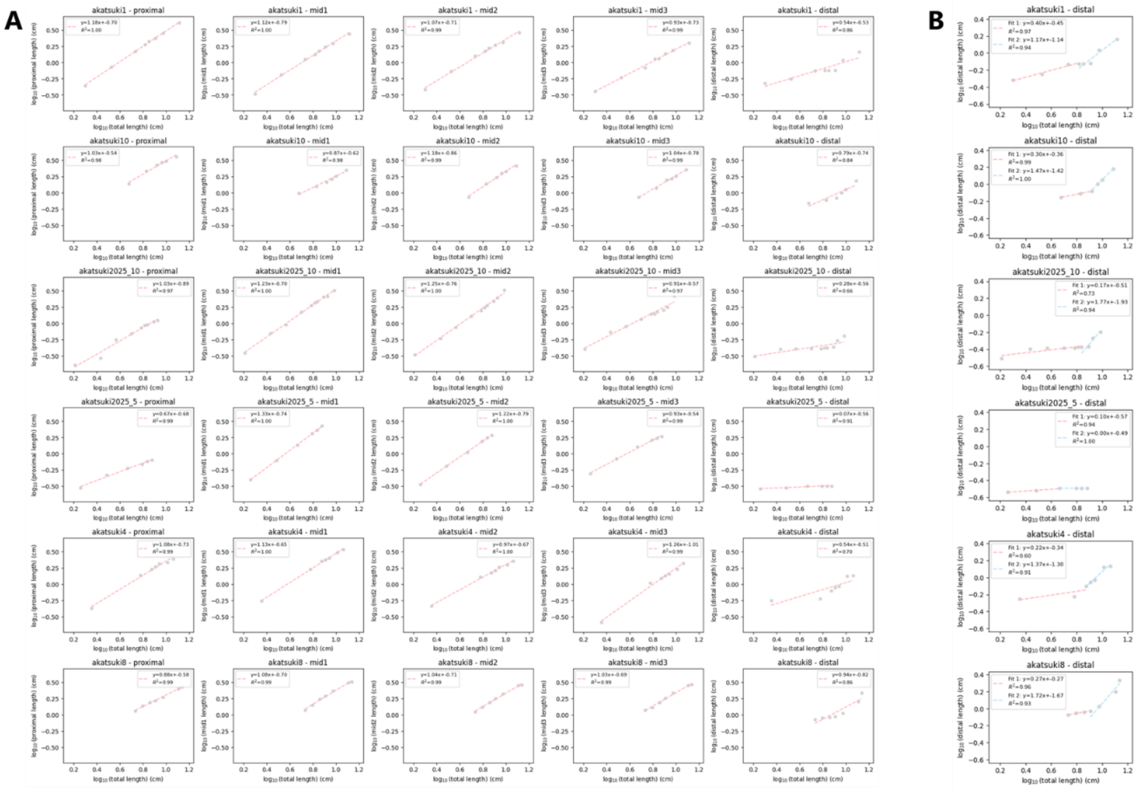


Figure S7. Allometric analysis of spatial growth in peach ‘Akatsuki’. (A) Single linear regression applied to all regions. From left to right: proximal, mid1, mid2, mid3, and distal regions. (B) Piecewise linear regression applied only to the distal region.

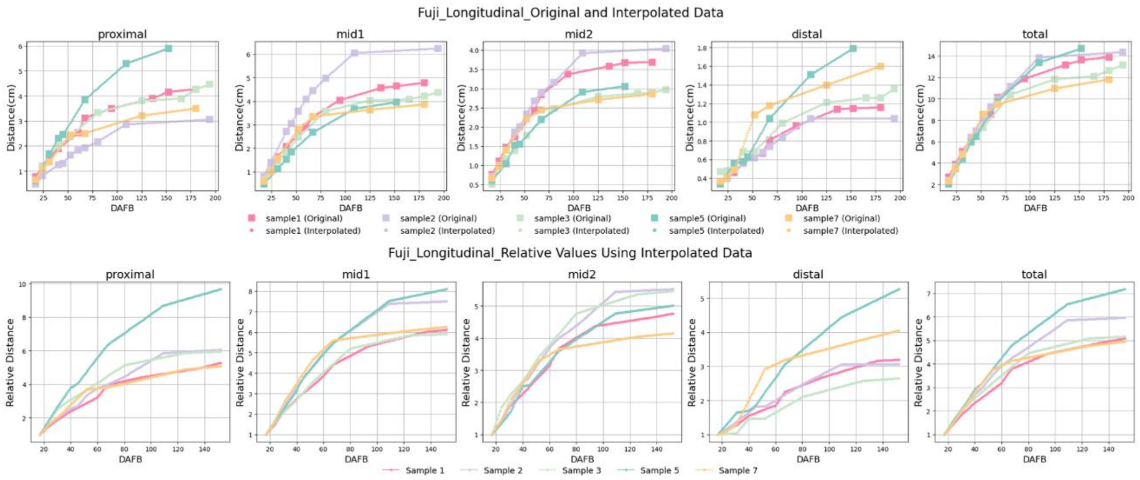


Figure S8. Daily growth curves obtained by spline interpolation for ‘Fuji’. Top: Graphs showing the raw (squares) and spline-interpolated (circles) daily distances between landmarks for each sample. Bottom: Relative distances for each sample, calculated by setting the latest initial day of 3D reconstruction among the samples as 1. From left to right: proximal, mid, mid2, distal, and total (sum of all regions).

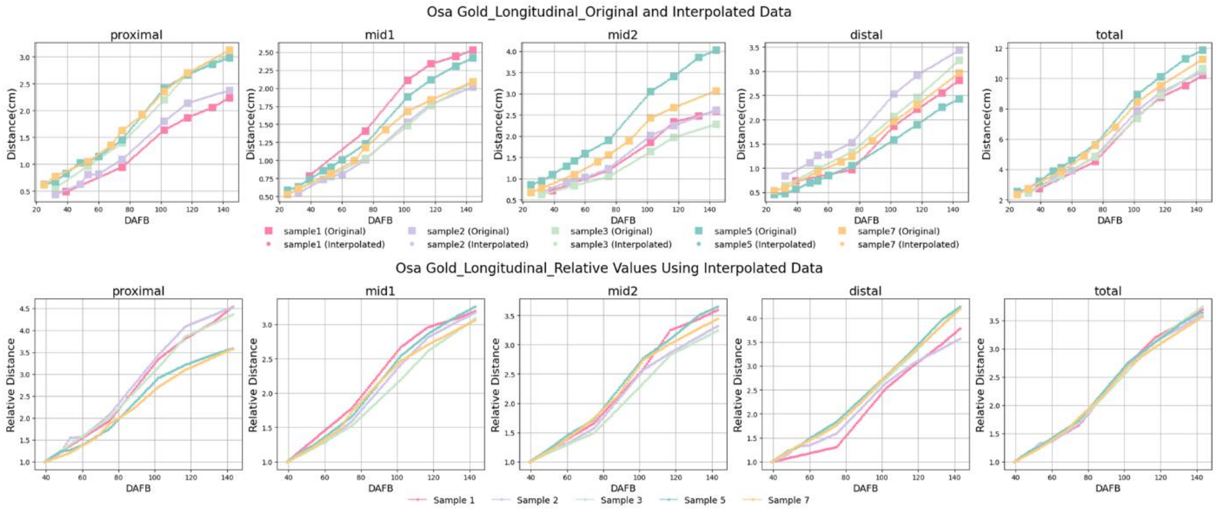


Figure S9. Daily growth curves obtained by spline interpolation for ‘Osa Gold’. Top: Graphs showing the raw (squares) and spline-interpolated (circles) daily distances between landmarks for each sample. Bottom: Relative distances for each sample, calculated by setting the latest initial day of 3D reconstruction among the samples as 1. From left to right: proximal, mid, mid2, distal, and total (sum of all regions).

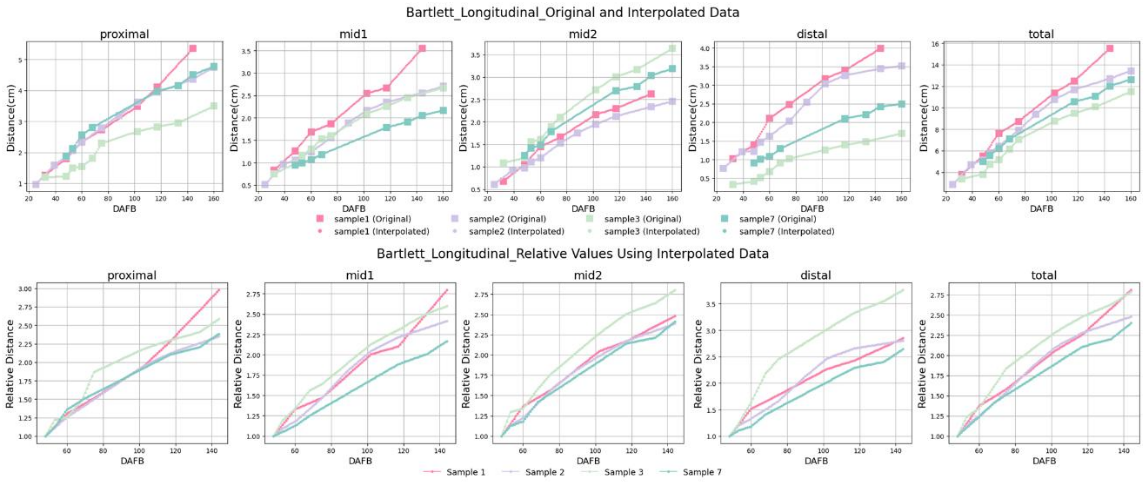


Figure S10. Daily growth curves obtained by spline interpolation for 'Bartlett'. Top: Graphs showing the raw (squares) and spline-interpolated (circles) daily distances between landmarks for each sample. Bottom: Relative distances for each sample, calculated by setting the latest initial day of 3D reconstruction among the samples as 1. From left to right: proximal, mid, mid2, distal, and total (sum of all regions).

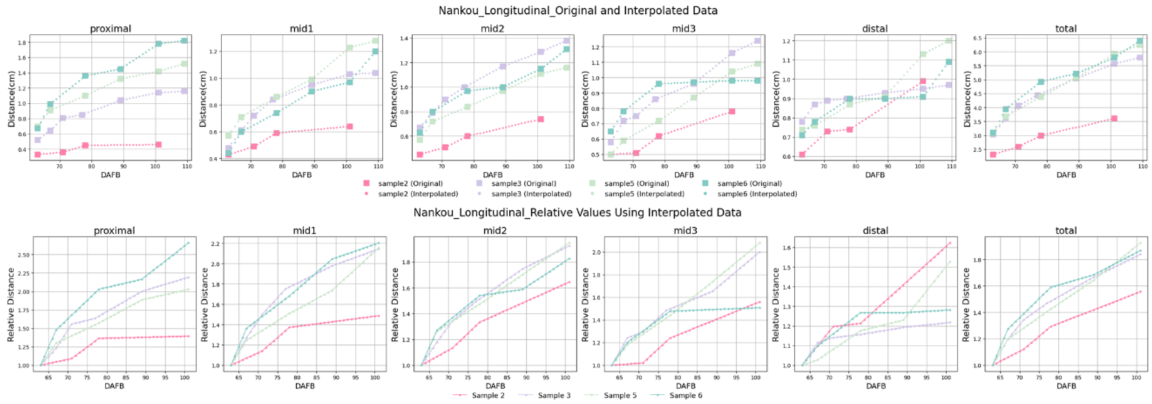


Figure S11. Daily growth curves obtained by spline interpolation for 'Nankou'. Top: Graphs showing the raw (squares) and spline-interpolated (circles) daily distances between landmarks for each sample. Bottom: Relative distances for each sample, calculated by setting the latest initial day of 3D reconstruction among the samples as 1. From left to right: proximal, mid, mid2, mid3, distal, and total (sum of all regions).

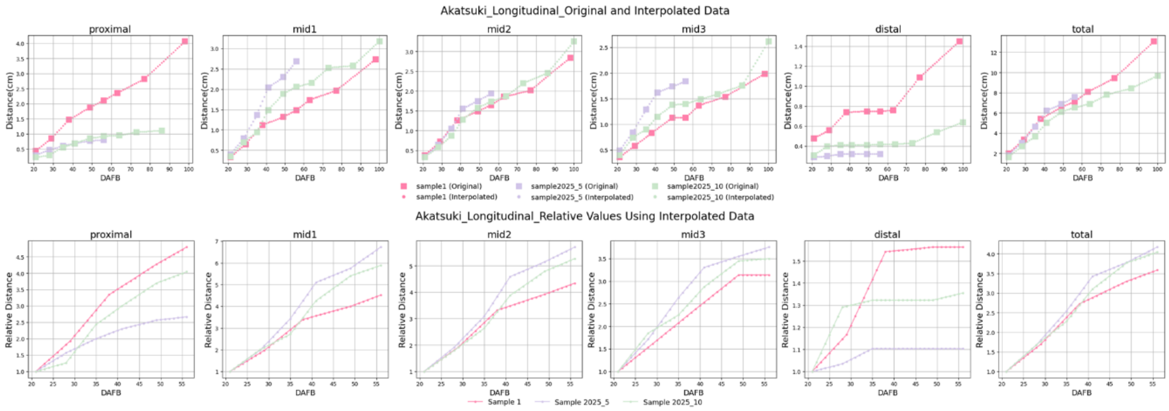


Figure S12. Daily growth curves obtained by spline interpolation for 'Akatsuki'. Top: Graphs showing the raw (squares) and spline-interpolated (circles) daily distances between landmarks for each sample. Bottom: Relative distances for each sample, calculated by setting the latest initial day of 3D reconstruction among the samples as 1. From left to right: proximal, mid, mid2, mid3, distal, and total (sum of all regions).
